# A robust mutation spectrum analysis framework for high-depth error-corrected sequencing data

**DOI:** 10.64898/2026.10.19.760182

**Authors:** Andrew Williams, Alper J Alcaraz, Annette E. Dodge, Danielle P.M. LeBlanc, Matthew J. Meier, Carole L. Yauk, Francesco Marchetti

## Abstract

Comparing mutational spectra across experimental groups requires inference at the level of the biological replicate. Error-corrected sequencing (ECS) generates hundreds to thousands of mutations per replicate, yet classical pooled G² tests treat individual mutations as independent observations and ignore replicate-level variability, inflating statistical significance (pseudo-replication). We present a replicate-aware Ridge-Penalized Dirichlet-Multinomial (RP-DMM) framework that models biological replicates directly and accounts for overdispersion. An L2 (ridge) penalty, with strength chosen by leave-one-out cross-validation, stabilizes spectrum estimates by shrinking weakly supported mutation categories toward a common baseline, and inference uses a parametric bootstrap. We also provide a complementary non-parametric rank-permutation test for sparse datasets.

In simulations, the pooled G² test became increasingly anti-conservative as mutation depth increased, with false-positive rates approaching 100% under the null hypothesis. RP-DMM maintained Type I error near the nominal 5% level and had substantially greater power than the rank-permutation test. Regularization was most consequential for high-dimensional spectra. False positive rates approached the nominal 5% level once the lower-mutation group contained approximately 300-500 mutations, depending on the number of biological replicates. A practical target of approximately 400 mutations in the lower-mutation group provided robust Type I error control across all scenarios examined; below this, the rank-permutation test performed robustly and is preferred. In Duplex Sequencing data from benzo[a]pyrene-exposed mice, RP-DMM distinguished exposure-associated mutational signatures from comparisons with higher mutation counts but null effect size, whereas the pooled G² test declared all comparisons significant. The cross-validated penalty decreased as mutation depth increased, indicating that regularization strength should be tuned to the data.

The framework is implemented in the MutSeqR R package.

## 1. INTRODUCTION

The statistical comparison of mutational spectra is essential for understanding DNA damage and repair across diverse biological systems and experimental conditions (Alexandrov et al., 2020). Historically, categorical approaches such as those established by Adams and Skopek (1987) and Piegorsch and Bailer (1994) have been widely used, relying on contingency table-based analyses suited to datasets containing modest numbers of mutant isolates per group.

With the advent of high-throughput and error-corrected sequencing (ECS), such as Duplex Sequencing and other consensus sequencing approaches (Marchetti et al., 2023; Yauk et al., 2026), contemporary studies routinely generate hundreds to thousands of mutational calls per sample, often across multiple biological replicates (Salk et al., 2018; LeBlanc et al., 2022; Dodge et al., 2023). While this increased resolution enhances statistical power, it also introduces significant analytical challenges: classical multinomial methods, such as the *G^2^* or Chi-square tests, can become overpowered and yield spurious significance as the total number of informative duplex bases and consequently the number of mutation calls increases (Salk et al., 2018). This “Large N” effect can drive p-values toward zero even in the presence of negligible biological differences (Lin et al., 2013). Crucially, these classical methods fail to account for replicate-to-replicate variability (technical and biological overdispersion), which is a defining feature of modern genomic datasets.

The inflation of significance stems from the assumption, in classical contingency table tests, that individual mutations are independent and identically distributed observations. This framework treats the mutation as the primary random variable, leading to “pseudo-replication” when mutation counts are interpreted as independent biological evidence. Mutations are clustered within biological replicates and exhibit intra-sample correlation. As the total number of mutations (*N*) increases, classical tests can increasingly underestimate the true variance by ignoring the hierarchy of the experimental design. The biological replicate (sample) should therefore be treated as the primary random variable. The sample size is then defined by the number of replicates (*n*), rather than the total mutational count (*N*).

Hierarchical Dirichlet-multinomial (DM) models have recently been applied to mutational signature exposures in cancer genomics. Morrill Gavarró et al. (2025) developed a mixed-effects DM model with patient-level random effects and group-specific precision parameters to compare clonal and subclonal exposures in paired tumour samples. That framework is designed for cohorts of tens to hundreds of patients with paired observations and relies on asymptotic Wald tests. Genetic toxicology studies using ECS typically have far fewer independent replicates per group and, at higher spectral resolution, many sparse mutation categories, which motivates the regularized, bootstrap-based approach presented here.

We present a replicate-aware analytical framework implemented in the MutSeqR R package (Dodge et al., 2025). Our primary approach is a ridge-penalized Dirichlet-multinomial model (RP-DMM) that accounts for overdispersion by modelling replicate-level spectra directly. Its ridge (L2) penalty shrinks weakly supported mutation-category proportions toward a common baseline, reducing sensitivity to stochastic fluctuations in low-count categories. We also implemented a non-parametric rank-permutation test (Alvo et al., 2010) whose statistical power is not artificially inflated by high mutation counts. Key terminology used to distinguish between implementations of the RP-DMM framework throughout this paper is defined in Box 1. We demonstrate this framework using Duplex Sequencing mutation spectra from benzo[a]pyrene (BaP) exposed mice (LeBlanc et al., 2022). Together, RP-DMM and the rank-permutation test support biologically meaningful comparisons between experimental groups while guarding against inflated statistical significance caused by high mutation counts and technical noise.

### Box 1. Glossary of Key Terms

Regularization: The general process of adding a penalty term to a statistical model to stabilize parameter estimates and prevent overfitting, particularly when data are sparse or high-dimensional. In the RP-DMM framework, regularization shrinks weakly supported mutation category proportions toward a common baseline, reducing sensitivity to technical noise.

Unpenalized DM: The standard Dirichlet-Multinomial model applied without a regularization penalty. Used as a benchmark to evaluate the effect of regularization.

RP-DMM: The Ridge-Penalized Dirichlet-Multinomial model proposed in this paper. Parameters (mutation spectrum proportions and the concentration parameter) are jointly optimized under an L2 penalty, producing stable, regularized estimates. Statistical inference is performed via a parametric bootstrap using a constructed test statistic evaluated at the optimized parameters.

Concentration parameter (θ): An inverse measure of overdispersion in the Dirichlet-Multinomial framework. Higher values of θ indicate lower overdispersion (individual replicates cluster tightly around the group mean spectrum), while smaller values correspond to higher overdispersion (greater variability among replicates) (Chen & Li, 2013; Holmes et al., 2012).

L2 Penalty (Ridge Penalty): A regularization approach that discourages extreme parameter estimates by adding a penalty proportional to the squared magnitude of the model parameters to the objective function. In the RP-DMM framework, the L2 penalty shrinks mutation spectrum proportions toward a common baseline, stabilizing estimates for sparse mutation categories while retaining information from well-supported categories.

Fully Penalized RP-DMM: An implementation in which the L2 penalty is applied during both parameter estimation and evaluation of the constructed test statistic. This approach maximizes specificity by applying the strongest regularization throughout.

Hybrid RP-DMM: An implementation in which the L2 penalty is applied during parameter estimation only, while the constructed test statistic is evaluated using the unpenalized likelihood. Because the parameters are already stabilized by the penalty before evaluation, results are similar to the Fully Penalized implementation when all mutation categories are well represented (e.g., the 6-channel spectrum); the two can diverge in sparse, high-dimensional spectra.

## 2. METHODS

Individual mutational spectra were analyzed using publicly available Duplex Sequencing data from LeBlanc et al. (2022; SRA accession number PRJNA803048). This dataset comprises 24 independent biological replicates from mice exposed to olive oil (controls) or increasing doses of BaP dissolved in olive oil. The 6-channel single-base substitution spectra were used as an example to compare the replicate-aware DMM approach and rank-permutation test with the classical pooled *G^2^* test. For each pairwise comparison, results from RP-DMM and the rank-permutation test were compared with those from the classical pooled *G²* test.

### Classical pooled *G²* approach

The traditional mutation spectrum analysis uses a classical pooled *G^2^* likelihood-ratio test (Piegorsch and Bailer, 1994). For each group comparison, mutational spectra were first represented as mutation-category-by-replicate count matrices with *K* mutation categories, where *K* = 6 for the 6-channel spectrum analyzed here. For the pooled *G^2^*test, mutation counts were then summed across biological replicates within each exposure group to generate a pooled contingency table of mutation categories by group. The *G^2^* statistic was calculated from the observed and expected category counts under the null hypothesis that the two groups shared the same mutation spectrum. This approach treats individual mutation calls as independent observations and does not model replicate-to-replicate variability (overdispersion). It therefore provides a useful benchmark for evaluating how classical contingency table methods behave when applied to datasets with large numbers of mutation calls.

### Ridge-Penalized Dirichlet-Multinomial Model (RP-DMM)

Using the same mutation-category-by-replicate count matrices, we modeled replicate-level mutation spectra directly. Consider an experiment with *G* groups containing multiple biological replicates. For replicate *j* in group *i, x_ijk_* denotes the number of observed mutations of category *k*. The total mutation count in replicate *j* in group *i* is denoted by *N_ij_* = ∑*_k_ x_ijk_*. Each group *i* is characterized by a mean spectrum probability vector *p_i_* = (p*_i_*_1,_…,*p_i,K_*), the expected mutation-category proportions across replicates from that group, where 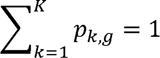.

To account for replicate-to-replicate variability (overdispersion), mutation counts for each sample were modeled as a draw from a Dirichlet-multinomial (DM) distribution (Mosimann, 1962; Rønning, 1989) with a group-specific concentration parameter. Related DM models with group-specific precision parameters and patient-level random effects have been used to compare mutational signature exposures between groups of tumours (Morrill Gavarró et al., 2025).

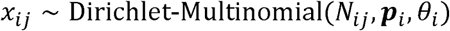

The DM distribution can be viewed as a two-stage model. Each replicate *j* in group *i* has its own underlying spectrum *π_ij_*, a vector of mutation-category probabilities that sum to 1, which varies around the group mean spectrum *p* according to a Dirichlet distribution with concentration parameter *θ*:*π_ij_* ∼ *Dirichlet(θp)*. Given *π_ij_*, the observed counts are multinomial: *x* | *π_ij_* ∼ *Multinominal ( N,π_ij_*). The DM probability mass function is obtained by integrating (marginalizing) the multinomial PMF over the unobserved *π_ij_*, weighted by the Dirichlet density, across all possible spectra (the (K − 1)-simplex). The result is a distribution with the same mean as the multinomial but greater variance, which is how the model captures replicate-to-replicate variability.

The concentration parameter *θ* is the inverse of overdispersion. Higher values indicate lower overdispersion, with replicates clustering tightly around *p*, while lower values indicate greater overdispersion (Holmes et al., 2012; Chen & Li, 2013). Because θ is estimated separately for each group, we also used it as a measure of within-group replicate consistency: low values may indicate technical outliers or unexpected biological heterogeneity. To stabilize the model when mutation counts are large or spectra are high-dimensional (e.g., the 96-base spectrum), we implemented a Ridge-penalized (*L2*) log-likelihood approach (Hoerl & Kennard, 1970). The penalized log-likelihood for group *i* is defined as:

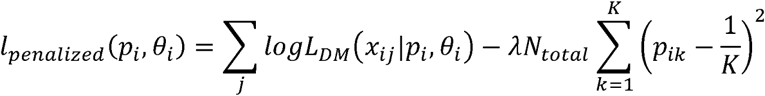

Where *logL_DM_* is the standard Dirichlet-Multinomial log-likelihood, *λ* is the regularization parameter, and *N_total_* is the total mutation count across the group. The penalty term is explicitly scaled by *N_total_* to ensure that regularization strength remains proportional to total number of mutations. This prevents the high statistical power of large datasets from magnifying minor technical fluctuations into spurious significance by shrinking the estimated spectrum toward a uniform distribution (1/K), which serves as a non-informative null prior in the absence of strong evidence for specific mutation types.

### Model Comparison and Inference

Statistical significance was assessed using a Likelihood Ratio Test (LRT) framework designed to maximize sensitivity while maintaining robustness. For each comparison, we evaluated two hypotheses: a null hypothesis (*H_0_*), assuming all groups share a common mutation spectrum and concentration parameter, and an alternative hypothesis (*H_A_*), allowing for group-specific parameters. The concentration parameter (θ) is treated as a nuisance parameter in this framework; while it is estimated under both hypotheses to account for overdispersion, the primary inferential target is the mutation spectrum (**p**). Consequently, a significant result reflects a composite difference between groups that may include differences in spectral composition, replicate consistency, or both.

Model parameters for both hypotheses were estimated by maximizing the ridge-penalized log-likelihood at a fixed value of λ, applied identically under HL (pooled data), under H_A (group-specific fits) and in every bootstrap replicate, to ensure stable estimates across mutation contexts. In the simulations, λ was fixed at 1.0. For the BaP analysis, λ was chosen beforehand by leave-one-out cross-validation (LOOCV) over biological replicates, performed separately within each dose group (select_optimal_lambda_joint, MutSeqR). For each candidate λ (0.001, 0.01, 0.05, 0.1, 0.5, 1, 2, 5), the spectrum proportions and concentration parameter were jointly estimated on the retained replicates, and the unpenalized log-likelihood of the held-out replicate was evaluated; the λ with the highest summed held-out log-likelihood was selected. Because the cross-validation surfaces were flat at low λ (see Results), a single value, λ = 0.001, was used for all pairwise comparisons. The constructed test statistic was defined as Λ = 2(*l_alt_* -*l_null_*), where l is the unpenalized log-likelihood (λ = 0) evaluated at the previously estimated regularized parameters.

The hybrid approach uses a constructed statistic rather than a classical LRT. Because the Ridge penalty alters the distribution of the test statistic, standard asymptotic distributions, such as the *χ*^2^distribution, were not used. Instead, p-values were obtained empirically via a parametric bootstrap (1000 replicates). For each bootstrap replicate, data were simulated using the shared parameters derived under the null hypothesis; parameters were then re-estimated using the ridge penalty, and the test statistic was evaluated using the unpenalized likelihood (λ=0). The p-value was calculated as the proportion of bootstrap replicates where the simulated hybrid test statistic exceeded the observed value (Efron & Tibshirani, 1993).

The study included six pre-specified pairwise comparisons for which raw p-values are reported. For studies involving larger numbers of comparisons, RP-DMM p-values can be adjusted using standard multiple testing correction methods (e.g., Holm-Sidak).

### Hybrid and Fully Penalized Test Statistics

Parameters under *H*_0_ and *H_A_* were always estimated by maximizing the ridge-penalized log-likelihood. We considered two ways of evaluating the test statistic, as Λ = 2(*l_alt_* - l*_null_*), at these estimates. In the Hybrid implementation, l is the unpenalized log-likelihood (λ = 0). In the Fully Penalized implementation, l is the penalized log-likelihood at the same λ. The implementations therefore share all parameter estimates and differ only in how the statistic is evaluated. The difference between the two statistics can be expressed analytically:

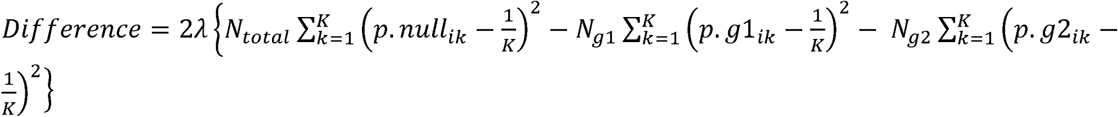

where *g*_1_ denotes group 1 and *g*_2_ denotes group 2.

This difference is zero, and the two statistics are equivalent, when:

• λ = 0. Both statistics reduce to the standard unpenalized likelihood ratio statistic.

• The count-weighted penalty under *H_A_* equals the penalty under *H_0_*(*N*₁*P*(p̂_1_) + *N*₂*P*(p̂_0_) = *N* · *P* (*p̂*0). This holds approximately when the two group spectra are similar to each other, as under a true null.

The statistics diverge when the groups’ spectra differ. When the null spectrum is approximately the count-weighted mean of the group spectra, the combined group penalties exceed the null penalty by (*N*₁*N*₂/*N*) *Σ*(*p̂*₁- *p̂*₂)², because pooling can only reduce the squared distance from the uniform baseline. Consequently, Λ_*FP* - Λ_*Hybrid* ≈ −2*λ*.(*N*₁*N*₂/*N*) *Σ*(*p̂*₁- *p̂*₂): the Fully Penalized statistic is the smaller one, and the gap grows with λ, with the mutation counts, and with the squared distance between the two group spectra. The Fully Penalized statistic is thus suppressed in proportion to the size of the spectral difference, so the Fully Penalized test can lose power to detect real differences. The Hybrid approach avoids this by evaluating significance with the unpenalized likelihood, after the ridge penalty has stabilized the parameter estimates.

Because the bootstrap recomputes both statistics on every simulated null dataset, the penalty term enters the null distribution as well as the observed statistic, and p-values for both implementations remain valid under H&#25A1; regardless of λ (Supplemental Results).

### Non-parametric Rank-Based Comparison

We also implemented a non-parametric rank-permutation test based on the multivariate ranking logic described by Alvo et al. (2010). Within each sample, the *K* mutation categories were ranked from most to least frequent (rank 1 = highest count, rank *K* = lowest), with tied categories, including those with zero counts, assigned the average of the ranks they span (e.g., two categories tied for the 3rd and 4th highest count both receive rank 3.5).

A test statistic, *D_rank_*, was defined as the Euclidean distance between the mean rank vectors of the comparison groups:

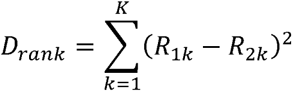

where *R_ik_* represents the mean rank of mutation category *k* in group *i*. Statistical significance was determined via a permutation test (1,000 iterations) by randomly shuffling group labels and recomputing *D_rank_* to generate a null distribution. This approach provides a total number of mutations-invariant sensitivity analysis that focuses purely on the relative ordering of mutation types within the spectra.

### Simulation Study

We evaluated the performance and Type I error rates of the classical pooled G² test, the rank-permutation test, and both RP-DMM implementations (Hybrid and Fully Penalized) using simulated mutational data. To create a true null scenario, in which no biological difference between groups was present, and with realistic overdispersion, spectra for two groups (4 replicates each) were generated from the same Dirichlet-multinomial distribution 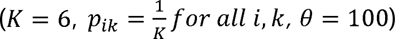. Under this design, any statistically significant result represents a false positive. We then assessed the impact of the “Large N” effect by varying the total number of mutations across four orders of magnitude (10, 100, 1,000, and 10,000 mutations per sample). For each scenario,1000 independent simulations were performed, and the Type I error was calculated as the proportion of simulations yielding significant results (p < 0.05). A sample size of 4 per group is consistent with the recommended minimum sample size for Duplex Sequencing mutagenicity studies (Esina et al., 2024, Williams et al., 2026).

We also evaluated the performance and Type I error rates of the G² test, the rank-permutation test, and Hybrid Implementation using simulated mutational data for the 96 trinucleotide spectrum. To create a true null scenario in which no difference in mutational spectrum was present between groups, spectra for two groups were generated from the same Dirichlet-multinomial distribution with (K=96) mutation categories and θ = 100. Under this design, statistically significant results represent false positives. We first evaluated Type I error under matched mutation burdens between groups (1×), and then assessed the effect of unequal mutation burdens while maintaining identical underlying spectrum proportions by allowing the treated group to contain 1.25-, 1.5-, or 2-fold more mutations than the control group. Mutation depth was varied across a range of per-sample mutation counts, and the number of replicates per group was varied from 4 to 6. For each combination of mutation depth, replicate number, and relative mutation burden, 1,000 independent simulations were performed. Type I error was calculated as the proportion of valid simulations yielding a significant result (p<0.05). Expected mutation counts per category were also calculated to characterize the relationship between total mutation depth and the sparsity of the 96 trinucleotide spectrum. These simulations were designed to identify practical mutation-depth requirements for reliable application of RP-DMM to high-dimensional mutational spectra.

### Application to Benzo[a]pyrene Duplex Sequencing data

We analyzed mutation spectra from mice exposed to various doses of BaP sequenced using Duplex Sequencing (LeBlanc et al., 2022). For each pairwise comparison across the four exposure groups (control, low, mid, and high), we applied three analytical approaches: the classical pooled G² test, the rank-permutation test, and the RP-DMM framework. Results from each approach were compared to evaluate the impact of replicate-aware modeling on the detection of exposure-associated spectral shifts.

### Concentration Parameter Interpretation

For each comparison, we report the maximum likelihood estimate of the Dirichlet-multinomial concentration parameter, θ. As mentioned above, in this framework θ serves as an inverse measure of overdispersion, quantifying replicate-to-replicate consistency by measuring how closely individual samples cluster around the group mean spectrum. High θ values indicate minimal overdispersion and strong consistency among replicates, while lower θ values correspond to greater overdispersion, reflecting higher variability among replicates and a less coherent group spectrum. We therefore report θ as a summary of the within-group replicate consistency alongside the comparative results.

## 3. RESULTS

### Validation via Simulation: Mitigation of the “pseudo-replication” Effect

We compared the Type I error rates of the classical pooled G² test, the rank-permutation test, and both RP-DMM implementations (Hybrid and Fully Penalized) using simulated mutational data. Under a true null hypothesis, in which the underlying mutation spectra are identical between groups, the empirical false-positive rate (FPR), corresponding to the estimated Type I error rate, should be approximately 5% at a nominal significance level of α = 0.05.

For the 6-channel spectrum (K = 6), the classical G^2^ test exhibited near-nominal Type I error at very low mutation counts (N = 10, FPR = 7%), but its false-positive rate increased as mutation depth increased. At 100 mutations per sample, the G^2^ test exhibited a 35% FPR, increasing to 96% at N = 1,000 and 100% at N = 10,000 mutations per sample (Figure 1). In contrast, the rank-permutation test, the Fully Penalized RP-DMM, and the Hybrid RP-DMM all maintained stable Type I error rates between 2% and 7% across the entire tested range. The near-identical performance of the two RP-DMM implementations is consistent with the theoretical expectation that joint optimization under the ridge penalty stabilizes both methods equivalently in the low-dimensional 6-channel spectral context, where all mutation categories are well-represented at all tested depths. These results indicate that by accounting for replicate-to-replicate variability, the RP-DMM framework effectively distinguishes technical noise from biological signal, providing a reliable alternative to traditional contingency table approaches that fail under pseudo-replication conditions.

**Figure 1.**
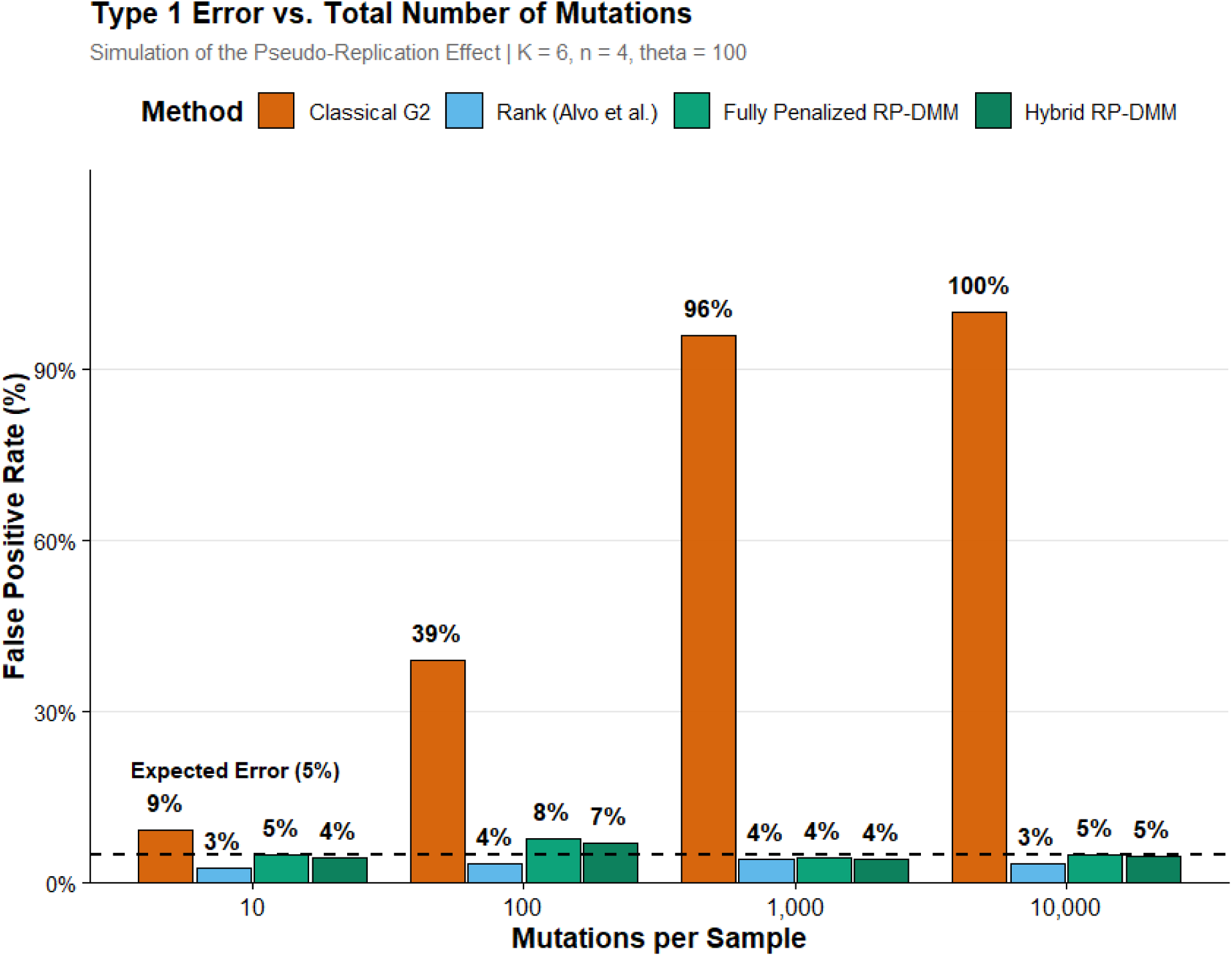
Impact of total mutation count on the false positive rate of four analytical approaches under a true null hypothesis. Simulations were performed under a null scenario in which both groups were generated from identical underlying mutation spectra (K = 6 categories, θ = 100, n = 4 replicates per group) across four mutation depths (10, 100, 1,000, and 10,000 mutations per sample; 1000 simulations per depth). Model parameters for both RP-DMM implementations were jointly optimized under the ridge penalty (λ = 1.0), with the Hybrid RP-DMM (dark green) evaluating the constructed test statistic using the unpenalized likelihood and the Fully Penalized RP-DMM (light green) using the penalized likelihood. Statistical inference for both RP-DMM implementations was performed using 1000 parametric bootstrap replicates. The dashed horizontal line indicates the nominal 5% significance threshold. As mutation depth increased, the classical pooled G² test (orange) exhibited increasing false positive rates, reaching 35%, 96%, and 100% at 100, 1,000, and 10,000 mutations per sample, respectively. In contrast, the rank-permutation test (light blue), Fully Penalized RP-DMM (light green), and Hybrid RP-DMM (dark green) maintained false positive rates between approximately 3% and 7% across all mutation depths.

For the 96 trinucleotide spectrum (K = 96), the Type I error behavior of the RP-DMM framework differed substantially from that observed for the 6-channel spectrum, highlighting the greater mutation-depth requirements imposed by the higher-dimensional spectrum (Supplementary Figure 1). At extremely low mutation depths mutations per sample (*N_i_* = ∑_k_ *x_ik_* = 10); group total N = 40), both RP-DMM implementations exhibited severely inflated Type I error rates (99%), reflecting the extreme sparsity of the 96 trinucleotide spectrum and the resulting difficulty of estimating the Dirichlet-multinomial parameters. The ridge penalty directly stabilizes the spectrum proportions (**p**) by shrinking them toward the uniform baseline, but does not directly penalize θ. Consequently, at very low mutation counts, estimation of θ can become poorly constrained, potentially contributing to instability in the penalized likelihood-ratio test. This instability was attenuated as mutation depth increased, with Type I error approaching nominal levels only at substantially greater mutation depths in the initial simulations. These results demonstrate that substantially greater mutation depth is required for reliable inference with the 96 trinucleotide spectrum than with the lower-dimensional 6-channel spectrum.

To further characterize the minimum mutation depth required for reliable inference, additional simulations were performed across a range of mutation counts, numbers of replicates, and relative mutation burdens between groups (Supplemental Table 2). These simulations indicate that the classical guideline of five expected observations per category (Cochran, 1954), corresponding to approximately 480 mutations across the 96 trinucleotide spectrum, provides a reasonable approximation for the minimum mutation depth required by the RP-DMM framework. False positive rates remained substantially elevated at lower mutation counts, but approached the nominal 5% level once the lower-mutation group contained approximately 300–500 mutations, with similar performance observed when the comparison group had 1.25- to 2-fold greater mutation burden. Increasing the number of replicates allowed reliable Type I error control at somewhat lower mutation counts, whereas substantially greater mutation depths provided little additional improvement. These results support a practical minimum target of approximately 300–500 mutations in the lower-mutation group for application of RP-DMM to 96-channel trinucleotide mutational spectra, with greater mutation counts preferred where feasible. In contrast, the rank-permutation test maintained near-nominal Type I error rates (3–5%) across the entire range of mutation depths examined in the 96-base spectral context, including the sparsest scenario (group total = 40; Supplementary Figure 1). The classical pooled G² test exhibited near-nominal Type I error only at the lowest mutation depth, not because the method is appropriate for sparse data, but because the limited number of mutations severely restricts its ability to detect any differences, including spurious ones. As mutation depth increased, however, the G² test rapidly became anti-conservative, with false positive rates approaching 100% (Supplementary Figure 1), consistent with its behavior in the 6-base spectral context (Figure 1).

These findings support a mutational-depth guideline for RP-DMM. Under the simulation conditions considered here, reliable RP-DMM inference for the 96-channel trinucleotide spectrum required approximately 400 total mutations in the lowest mutational burden group, corresponding to roughly 100 mutations per sample when four biological replicates are available. These simulations suggest a practical guideline of approximately 4-5 × K total mutations in the lower-mutational-burden group., corresponding to roughly 300-500 mutations for a 96-channel spectrum. This recommendation is empirical and was derived from simulations assuming a uniform null mutation spectrum. Additional simulations are needed to determine whether different underlying mutational signatures require higher or lower mutational depths. Below this practical target, the rank-permutation test maintained nominal Type I error under sparse-data conditions.

The increasing false-positive rate of the classical G² test reflects its treatment of each mutational call as an independent observation. The classical G² test ignores the clustering of mutations within biological replicates, causing statistical significance to increase with mutation depth rather than biological effect size. In contrast, the replicate-aware RP-DMM framework and the rank-permutation test maintained nominal Type I error across the tested mutation depths in the 6-channel spectral context. The near-identical performance of the Hybrid and Fully Penalized RP-DMM implementations was observed in this low-dimensional setting, and their Type I error behavior remained closely matched even in the higher-dimensional 96-channel context (Supplementary Figure 1), despite the two implementations later diverging in statistical power under sparse, high-dimensional conditions (see below).

### Sensitivity and Limit of Detection

The classical pooled G² test was excluded from sensitivity and power analyses because its inflated Type I error rates under the null hypothesis rendered power estimates uninterpretable. To characterize sensitivity, we simulated increasing perturbations in the C>A mutation category while maintaining a total of 1,000 mutations per biological replicate and four replicates per group (Figure 2). The Hybrid and Fully Penalized RP-DMM implementations exhibited nearly identical performance across the tested range, indicating that evaluation using the penalized or unpenalized likelihood had minimal impact once model parameters were jointly regularized.

**Figure 2.**
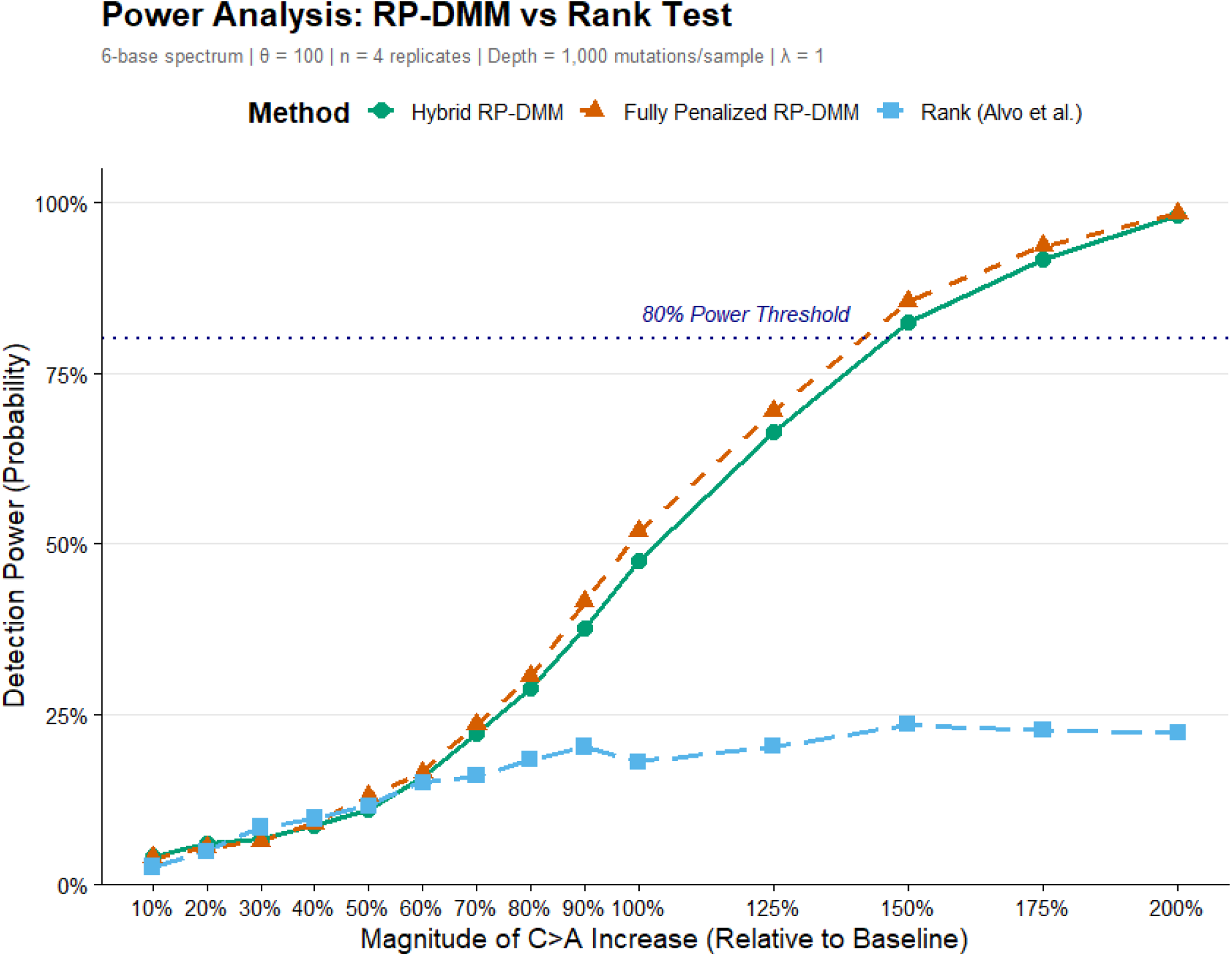
**Statistical power of the RP-DMM and rank-permutation tests under increasing spectral differences.**

Both RP-DMM implementations achieved approximately 80% statistical power at a 75% relative increase in the target mutation category, whereas the rank-permutation test remained substantially underpowered and did not exceed 30% power across the evaluated effect sizes.

The superior performance of RP-DMM reflects its use of mutation-count magnitudes, which are discarded during rank transformation. Together, these findings demonstrate that replicate-aware modeling provides substantially greater sensitivity for detecting mutational spectrum shifts while maintaining appropriate Type I error control.

Simulations were performed using 1,000 mutations per biological replicate and four biological replicates per group. The underlying mutation spectrum was modified by increasing the relative weight of the C>A mutation category across a range of effect sizes, with 1000 independent simulations performed at each effect size. The Hybrid RP-DMM (green, solid line) and Fully Penalized RP-DMM (orange, dashed line) used 500 parametric bootstrap replicates, whereas the rank-permutation test (blue, dashed line) used 500 permutations. Statistical power was calculated as the proportion of simulations yielding p < 0.05.

### Application to Mutational Spectra in Benzo[a]pyrene-Exposed Mice

We applied the replicate-aware RP-DMM framework to compare mutation spectra at four BaP exposure levels (control, low, mid, and high) in the LeBlanc et al. (2022) 6-channel single-base substitution dataset. The analysis included determining the appropriate regularization parameter, checking for consistency of the replicated data, and conducting the pairwise comparisons. We then compared the RP_DMM results with those from the classical pooled G^2^ test and rank statistic.

The regularization parameter (λ) was optimized independently for each dose group via leave-one-out cross-validation (LOOCV), with results summarized in Table1. The optimal λ decreased monotonically with increasing BaP dose and consequently with increasing total mutation counts: control (λ = 0.1, ∼130 mutations/sample), low (λ = 0.01, ∼260 mutations/sample), and mid and high dose groups (λ = 0.001, ∼350 and ∼740 mutations/sample, respectively).

This monotonic decrease in the optimal regularization parameter (λ) with increasing sample mutation depth is consistent with the role of the ridge penalty in the stabilizing mutation spectrum estimates. L2 regularization stabilizes spectrum estimates when mutation categories have zero or near-zero counts and are weakly supported by the data. In low-count datasets, such sparsity is common, and without regularization, stochastic fluctuations in sparse categories can drive spurious differences between groups. As the total number of mutations per sample increases, all mutation categories become sufficiently represented and require less regularization. Stronger penalization toward the uniform distribution can then suppress the strong, non-uniform BaP mutational signature that is well-supported by the data.

Examination of the cross-validation likelihood surfaces revealed that the curves were notably flat across the low-λ region (λ = 0.001 to λ = 0.1) for all groups. For the control group, the difference in predictive log-likelihood between the optimal λ = 0.1 and the most conservative value of λ = 0.001 was only 0.207 units, a negligible loss in predictive performance relative to the steep drop of 10.0 units observed when λ exceeded 0.1. Similarly, for the low dose group, the difference between λ = 0.01 (optimal) and λ = 0.001 was only 0.010 units. In contrast, all groups exhibited rapid deterioration in predictive performance for λ ≥ 0.5, confirming that over-regularization is the primary risk to avoid.

The flat likelihood surfaces in the low-λ region indicate that a single global value of λ = 0.001 provides near-optimal predictive performance across all dose groups. For the mid and high dose groups, λ = 0.001 is the CV-optimal value. For the control and low dose groups, it represents a negligible departure from their respective optima and avoids stronger regularization of the higher-dose groups. The global value was therefore selected to provide limited regularization across groups without substantially reducing predictive performance.

Accordingly, λ = 0.001 was applied uniformly across all pairwise comparisons. This data-driven selection ensures that results are robust to the specific choice of penalty parameter, as confirmed by the flat likelihood surfaces observed in the low- λ region for all groups.

### Replicate Quality Report

Prior to pairwise statistical testing, the concentration parameter (θ) was estimated independently for each dose group to characterize within-group replicate consistency (Table 2). Because higher BaP exposure is expected to produce a stronger, more dominant mutational signature that increasingly outweighs background technical and biological noise, we anticipated θ would increase monotonically with dose. The parameter summarizes within-group consistency but does not identify the source of variability, which may reflect technical artifacts, biological heterogeneity in treatment response, or stochastic variation in mutation accumulation.

**Table 1.** Leave-One-Out Cross-Validation Results for Regularization Parameter Selection.

| **Dose Group** | **Total Mutations** | **Optimal λ** | **CV Score<sup>1</sup><br>(Optimal λ)** | **CV Score<br>(λ = 0.001)** | **Flatness Cost<sup>2</sup>** |
|---|---|---|---|---|---|
| Control | ~ 800 | 0.100 | -1174.478 | -1174.685 | 0.207 |
| Low | ~ 1550 | 0.010 | -2254.590 | -2254.600 | 0.010 |
| Mid | ~ 2,100 | 0.001 | -2641.057 | -2641.057 | 0 |
| High | ~ 4,450 | 0.001 | -5013.816 | -5013.816 | 0 |

<sup>1</sup>CV scores represent the sum of held-out log-likelihoods evaluated at the unpenalized likelihood (λ = 0) after joint optimization of the mutation spectrum proportions (*p*) and concentration parameter (θ) under each candidate’s λ value. Higher (less negative) scores indicate better out-of-sample predictive performance.
<sup>2</sup>Flatness Cost: The difference in predictive log-likelihood between the group-specific optimal λ and the global baseline value of λ = 0.001. Values close to zero indicate that λ = 0.001 provides near-optimal predictive performance for that group.

**Table 2.** Group-Specific Concentration Parameter Estimates and Replicate Consistency (λ = 0.001).

| **Dose Group** | **Total Mutations** | **Estimated θ<sup>1</sup>** | **Variability** |
|---|---|---|---|
| Control | 769 | 279.6 | Low |
| Low | 1589 | 248.3 | Low |
| Mid | 2082 | 41,538<sup>2</sup> | Minimal (Near-perfect) |
| High | 4423 | 2235.3 | Minimal |

<sup>1</sup>Concentration parameters estimated by joint optimization of spectrum proportions (p) and concentration parameter (θ) under the ridge penalty (λ = 0.001).
<sup>2</sup>Near-perfect replicate consistency. The Dirichlet-Multinomial concentration parameter is unbounded above; this value indicates that between-replicate variation in spectral proportions is negligible, consistent with a dominant mutational signal overwhelming background noise at this dose level.

At λ = 0.001, θ estimates ranged from 248.3 in the low dose group to 41,538 in the mid dose group (Table 2). The near-perfect replicate consistency of the mid dose group reflects the emergence of a dominant, highly reproducible BaP mutational signature at this exposure level, where the strong C>A signal overwhelms background technical variability across all replicates. The high dose group also demonstrated high replicate consistency (θ = 2,235.3), consistent with a well-established mutational signature at the highest exposure level.

Two observations deviated from this expected monotonic increase in θ with dose. First, the Low dose group (θ = 248.3) exhibited slightly lower replicate consistency than the Control group (θ = 279.6), despite having approximately twice the total mutation count (∼1,589 vs. ∼769 mutations). This pattern may reflect the early emergence of the BaP mutational signature in some replicates but not others at the low dose level, introducing between-replicate biological heterogeneity in addition to background technical noise. Second, while the high dose group had the highest total mutation count (∼4,423 mutations), its θ estimate (2,235.3) was substantially lower than that of the mid dose group (41,538). This deviation may reflect genuine biological heterogeneity in mutational response at the highest dose. However, as noted above, θ alone cannot distinguish between these possibilities, and examination of individual replicate spectra is recommended when interpreting these results.

These θ estimates provide important context for the pairwise comparisons reported in Table 3. Groups with high θ values (e.g., Mid dose) exhibit tight replicate clustering, whereas groups with lower θ values, such as the Control and Low dose groups, exhibit greater within-group variability. These differences in replicate consistency should be considered when interpreting the pairwise comparisons. We therefore report θ alongside the p-values as a summary ofreplicate consistency.

**Table 3.** Pairwise mutation spectrum comparisons across benzo[a]pyrene dose groups using the classical pooled G^2^ test, the non-parametric rank-permutation test, and the Ridge-Penalized Dirichlet-Multinomial (RP-DMM) framework (λ = 0.001).

| Comparison | ***G*<sup>2</sup> p-value** | **Rank p-value** | ***θ*** | **Full RP-DMM p-value** | **Hybrid RP-DMM p-value** |
|---|---|---|---|---|---|
| Control vs Low | < 0.001 | 0.202 | 135.89 | 0.010 | 0.010 |
| Control vs Mid | < 0.001 | 0.028 | 99.08 | <0.001 | <0.001 |
| Control vs High | < 0.001 | 0.009 | 63.36 | <0.001 | <0.001 |
| Low vs Mid | < 0.001 | 0.143 | 280.79 | <0.001 | <0.001 |
| Low vs High | < 0.001 | 0.020 | 139.59 | <0.001 | <0.001 |
| Mid vs High | < 0.001 | 0.597 | 866.53 | <0.001 | <0.001 |

**Note:** G² p-values were obtained using the classical pooled likelihood-ratio test treating all mutation calls as independent observations. Rank p-values were obtained via a permutation test (1,000 permutations) of the Euclidean distance between group mean rank vectors (Alvo et al., 2010). RP-DMM p-values were obtained via parametric bootstrapping (500 replicates). For the Hybrid implementation, model parameters were jointly optimized under the ridge penalty (λ = 0.001) and the constructed test statistic was evaluated using the unpenalized likelihood; for the Fully Penalized implementation, the same jointly optimized parameters were used, but the test statistic was evaluated using the penalized likelihood at the same λ. The concentration parameter (θ) represents the shared Dirichlet-Multinomial concentration under the null hypothesis.

### Pairwise Mutation Spectrum Comparisons Across BaP Dose Groups

The classical pooled G^2^ test yielded significant p-values (p < 0.001) for all six pairwise comparisons (Table 3), including comparisons where the spectral differences were visually subtle. The G² test treats each mutation call as an independent observation and does not account for clustering within biological replicates, which can inflate statistical significance as mutation counts increase

In contrast, the RP-DMM framework provided a more differentiated and biologically interpretable set of results. For example, in the control vs. low dose comparison, the classical pooled G² test yielded a highly significant result (p < 0.001), whereas RP-DMM detected a more modest, though still statistically significant, spectral shift (p = 0.010), consistent with the visually subtle difference between these two groups. The concentration parameter (θ) varied considerably across comparisons, ranging from 63.36 for the control vs. high dose comparison to 866.53 for the mid vs. high dose comparison (Table 3). This variation reflects the degree of spectral coherence across replicates when both groups are pooled under the null hypothesis. Lower θ values indicate greater between-replicate variability in the pooled dataset, which may result from large spectral difference between the two groups being compared, while higher θ values indicate that the combined dataset remains relatively homogeneous, as expected for more similar group spectra.

Comparisons involving groups with pronounced spectral differences were correctly identified as significant by the RP-DMM framework. The control vs. mid (p < 0.001), control vs. high (p < 0.001), low vs. mid (p < 0.001), and low vs. high (p < 0.001) comparisons all yielded highly significant results, consistent with the well-established dose-dependent C>A mutational signature of BaP. The lowest θ value was observed for the control vs. high comparison (θ = 63.36), indicating greater variability when the two groups were combined under the null model.

The control vs. low dose comparison yielded a significant RP-DMM p-value (p = 0.010) despite the relatively subtle visual spectral differences between these groups (Figure 3A). The shared concentration parameter for this comparison (θ = 135.89) was similar to values observed for other comparisons involving the low dose group (e.g., low vs. high, θ = 139.59; Table 3). At λ = 0.001, RP-DMM detected a difference between the control and low dose spectra. The rank-permutation test did not detect a significant difference for this comparison (p = 0.202), indicating lower power than RP-DMM under these conditions. The difference between the two results reflects their use of different information: RP-DMM uses mutation-count magnitudes, whereas the rank-permutation test uses the relative ordering of mutation categories within replicates.

**Figure 3:**
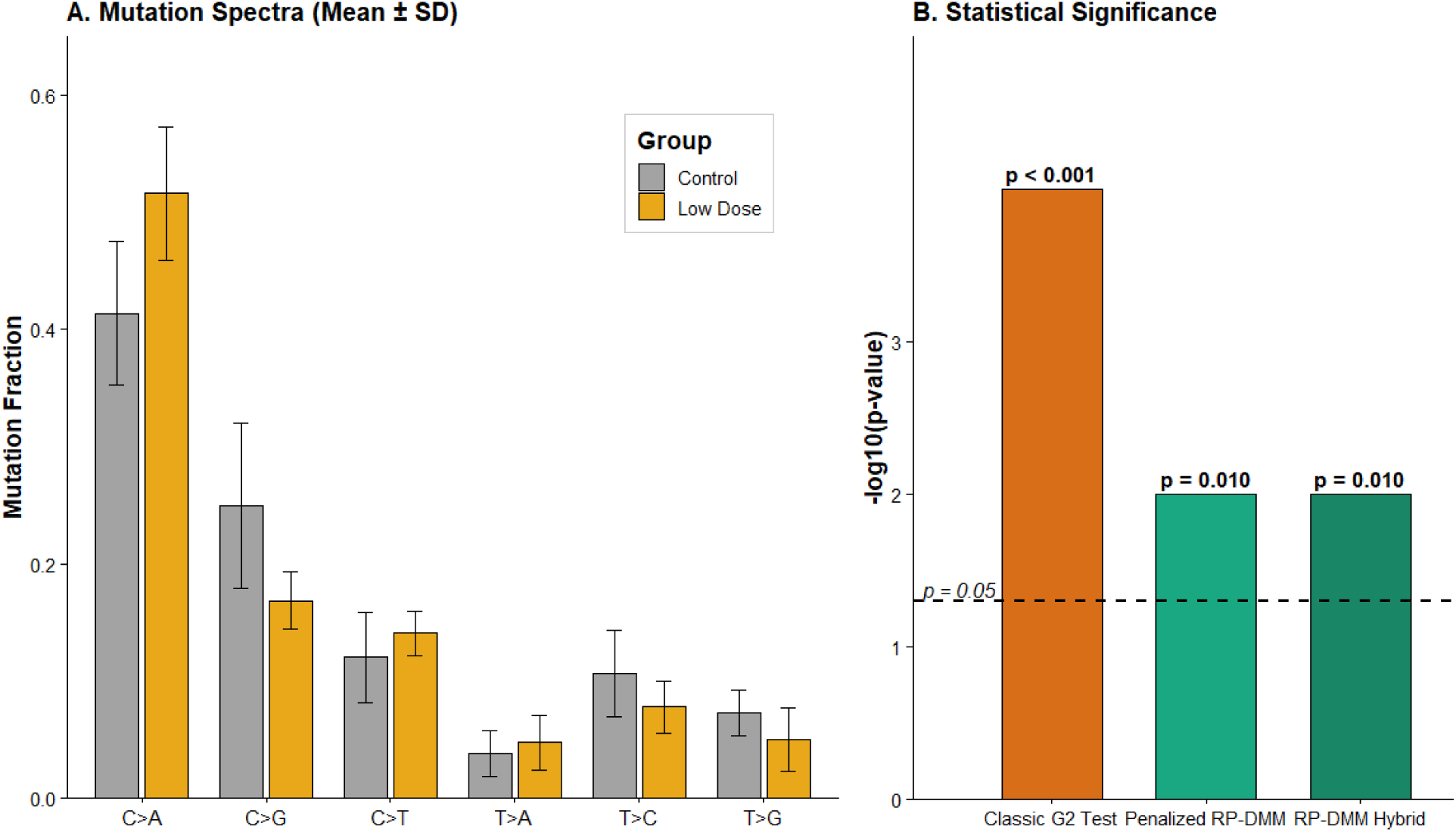
Comparison of mutation spectra between control and low dose benzo[a]pyrene-exposed mice. Panel A shows the mean (+/- SD) mutation fraction for each of the 6-channel single-base substitution categories in the control (grey) and low dose (gold) groups (n = 6 replicates per group). A modest increase in C>A mutations and decrease in C>G mutations is visible in the low dose group relative to control, consistent with early emergence of the BaP mutational signature. Panel B shows the statistical significance of the control vs. low dose comparison expressed as -log10(p-value) for three analytical approaches: the classical pooled G^2^ test (orange), the Fully Penalized RP-DMM (green), and the Hybrid RP-DMM (dark green).

The dashed horizontal line indicates the significance threshold (p = 0.05). The classical G^2^ test yielded a highly significant result (p < 0.001; Figure 3B). Both RP-DMM implementations, using joint optimization under the ridge penalty (λ = 0.001) with inference via parametric bootstrapping (500 replicates), yielded p = 0.010, reflecting a statistically detectable but subtle spectral shift consistent with early mutagenic exposure at the low dose level.

## 4. Discussion

Current approaches for mutation spectrum comparison, particularly the pooled G² framework, are widely used but can become anti-conservative in high mutation count contexts. Our implementation of the RP-DMM framework models replicate-level mutation spectra directly via a Dirichlet-multinomial distribution, accounting for overdispersion and replicate clustering that classical pooled tests ignore. A ridge penalty on the spectrum proportions further stabilizes estimates in weakly supported mutation categories, reducing sensitivity to sparse-category noise. Together, these two components ensure that statistical significance reflects reproducible spectral shifts rather than artifacts of high mutation counts or unmodeled replicate structure. In the simulation study, classical G² tests exhibited a near-100% false positive rate as the total number of mutations per sample increased, whereas the RP-DMM framework maintained statistical specificity near nominal levels across the entire tested range in the 6-channel spectral context (Figure 1); at higher spectral dimensionality (96 channels), nominal Type I error required a minimum mutation depth, as discussed above (Supplementary Figure 1). This ensures that flagged differences represent genuine biological shifts rather than pseudo-replication-driven statistical artifacts, within the depth ranges established here. A key strength of this framework is its positioning between overly sensitive contingency table approaches and strictly non-parametric methods. As demonstrated in our power analysis (Figure 2), the rank-permutation test (Alvo et al., 2010) provides a robust, mutation-count-invariant safeguard against spurious significance, but lacks the sensitivity required to detect subtle yet reproducible spectral shifts at the replicate numbers common in experimental toxicology. The RP-DMM framework preserves more biological information than rank-based methods by leveraging count magnitudes through the Dirichlet-multinomial distribution. The two methods are complementary rather than interchangeable: in the 96-channel simulations, RP-DMM’s Type I error control degraded sharply at low mutation depth (Supplementary Figure 1), where the rank-permutation test instead remained robust and is the preferred approach; at the higher mutation depths and lower spectral dimensionality typical of the BaP case study, RP-DMM’s greater sensitivity allowed it to detect differences the rank-permutation test missed, including the Control vs. Low (p = 0.202) and Mid vs. High (p = 0.597) comparisons.

The advantage of the RP-DMM framework at high spectral dimensionality is conditional on sufficient mutation depth. For 96-channel spectra, reliable Type I error control was generally achieved when the lower-mutational-burden group contained approximately 300-500 total mutations, with the exact requirement depending on replicate number. A target of approximately 400 mutations in the lower-burden group provides a practical and conservative recommendation for RP-DMM analyses (Supplementary Table 2). Above this depth, ridge regularization acts as a numerical anchor, shrinking sparse category estimates toward a common baseline and stabilizing the variance of individual trinucleotide contexts, giving RP-DMM greater sensitivity than rank-based methods once mutation counts are sufficient.

The target mutation depth may be achieved through additional biological replicates, increased sequencing depth, or both. Because statistical inference is driven primarily by the group with the lower mutation burden, studies should aim to ensure that the lowest-burden group contains sufficient mutations before applying RP-DMM in the 96-channel context. Importantly, these recommendations pertain to formal statistical inference and Type I error control and should not be interpreted as implying that mutational signatures cannot be estimated or described at lower mutation counts.

This depth recommendation has practical implications. In a survey of 28 vehicle-control Duplex Sequencing datasets from four published mutagenicity studies, total mutation counts per group ranged from 139 to 440 mutations (median = 206). Seven of the 28 datasets (25%) contained at least 300 mutations, whereas only three datasets exceeded 400 mutations. These observations indicate that many contemporary Duplex Sequencing vehicle-control datasets may operate near or below the mutation depths associated with reliable Type I error control in the present simulations (Williams et al., 2026). Consequently, studies intending to apply RP-DMM to 96-channel spectra should consider mutation depth during experimental design and ensure adequate mutation counts in the group with the lowest mutational burden.

The difference between the Hybrid and Fully Penalized test statistics is determined analytically by the difference in ridge penalty terms under the null and alternative hypotheses. As λ approaches zero, both statistics converge to the standard unpenalized likelihood ratio statistic, confirming that the two approaches are nested within the classical LRT framework. The two statistics are also equivalent when the penalty terms under *H*_0_ and *H_A_* are approximately equal, that is, when the null spectrum and the group-specific spectra deviate similarly from the uniform baseline (1/K), which is most likely when all categories are well-represented. This theoretical framework is supported empirically by our simulation results. In the 6-channel spectral context, the penalty terms under *H*_0_ and *H_A_* were approximately equal at λ = 0.001, and the two implementations produced near-identical Type I error rates and detection power (Figures 1 and 2). In the 96-channel trinucleotide mutation context, however, where category sparsity increases penalty-induced attenuation of the Fully Penalized test statistic, a meaningful divergence in detection power was observed between the two implementations (Supplementary Figure 2), with the Hybrid approach demonstrating approximately 10–15% higher power across the tested range. When one group carries a strong non-uniform mutational signature, the group-specific penalty term under *H_A_* becomes substantially larger than the null penalty term under *H*_0_ suppressing the Fully Penalized test statistic in proportion to the biological signal strength. The Hybrid approach avoids this additional penalization by evaluating significance using the unpenalized likelihood, allowing the biological signal to contribute fully to the test statistic once the parameter estimates have been stabilized by joint regularization. These findings demonstrate that the two approaches are effectively equivalent in low-dimensional contexts, whereas the Hybrid evaluation strategy provides a meaningful and empirically supported advantage in 96-channel and higher-dimensional mutation spectrum analyses, supporting its use for high-resolution signature comparisons in modern genomic studies.

To validate the performance observed in our simulation analyses, we applied the RP-DMM framework alongside classical and non-parametric approaches to mutation spectra generated in a BaP case study. The classical pooled G^2^ test identified all pairwise comparisons as statistically significant (p < 0.001), including comparisons between groups with visually subtle spectral differences. The pooled G² test treats each mutation call as an independent observation and does not account for clustering within biological replicates. The RP-DMM p-values varied across comparisons and were reported with the corresponding concentration parameter. In genetic toxicology, mutational spectra are increasingly used to assess mutagenicity, compare exposure conditions, and inform mechanistic interpretation (Yauk et al., 2026).

In the BaP case study, the selection of the regularization parameter was determined from the leave-one-out cross-validation. The optimal λ decreased monotonically with increasing BaP dose (control: λ = 0.1, low: λ = 0.01, mid and high: λ = 0.001), alongside increasing mutation depth. At low mutation depths, sparse mutation categories require regularization to prevent stochastic fluctuations from driving spurious differences. At high mutation depths, all categories are well-represented by the data, and stronger regularization toward the uniform distribution can suppress the dominant BaP mutational signature. Critically, examination of the cross- validation likelihood surfaces revealed that the curves were notably flat across the low- λ region for all groups, with the difference in predictive log-likelihood between the group-specific optimal λ and the global value of λ = 0.001 being negligible (control: 0.207 units; low: 0.010 units; mid and high: 0.000 units) (Supplemental Table 1.). This flatness justified the selection of a single global value of λ = 0.001 for all pairwise comparisons, providing near- optimal predictive performance across all dose groups simultaneously while ensuring that the strong BaP mutational signature in the higher dose groups was not suppressed by over-regularization.

Despite its robustness across a range of conditions, the RP-DMM framework has important limitations in ultra-sparse, high-dimensional settings. Simulations of the 96-channel trinucleotide spectrum indicated that reliable Type I error control was generally achieved when the lower-mutational-burden group contained approximately 300-500 total mutations, depending on the number of biological replicates, with approximately 400 mutations representing a practical target for routine applications. These recommendations should be regarded as empirical guidelines rather than universal requirements, as mutation-depth requirements may vary across underlying mutational spectra and experimental settings. The target mutation depth may be achieved through additional biological replicates, increased sequencing depth, or both. Because statistical inference was driven primarily by the group with the lower mutation burden, studies should aim to ensure that the lowest-burden group contains sufficient mutations before applying RP-DMM in the 96-channel context. Importantly, these recommendations pertain to formal statistical inference and Type I error control and should not be interpreted as implying that mutational signatures cannot be estimated or described at lower mutation counts. Below the mutation depths associated with reliable RP-DMM performance, the rank-permutation test maintained nominal Type I error rates and therefore provides a robust alternative for sparse mutational spectra.

The regularization parameter should be selected via the built-in cross-validation function rather than relying on a fixed default value, particularly for novel data types or spectral contexts not represented in the current study. The present analysis demonstrates that the optimal λ varied by total mutation depth (Supplementary Table 1), and that the choice of λ can materially affect the framework’s sensitivity to spectral differences: the same Control vs. Low comparison was non-significant at λ = 1 (p = 0.590) but significant at the cross-validated value, λ = 0.001 (p = 0.010; see Sensitivity Analysis). We would further expect optimal λ to depend on the strength of the underlying mutational signal, since a ridge penalty calibrated for weak, noisy spectra may over-shrink a strong, well-supported signature, though this was not directly tested here and warrants further investigation.

A further limitation concerns study design. Our framework complements mixed-effects DM models developed for cancer genomics (Morrill Gavarró et al., 2025), which model within-patient dependence in paired designs and report differential precision as a separate quantity from differential abundance. RP-DMM does not currently accommodate paired or repeated-measures designs, and because θ is treated as a nuisance parameter, a significant result may reflect differences in spectrum, in replicate consistency, or both. Extending RP-DMM to include random effects, or to test the spectrum and θ separately, is a natural direction for future work.

This study addresses the pseudo-replication problem inherent in pooled analysis. Under a true null hypothesis, the false-positive rate of the classical G² test increased with mutation depth, reaching 100% at the highest depth simulated, a finding that underscores the need for a shift in analytical standards. By introducing the replicate-aware RP-DMM framework with joint optimization and data-driven regularization, we align statistical inference with the actual experimental design of ECS studies, treating the biological replicate rather than the individual mutation as the primary unit of analysis. The RP-DMM framework offers a robust, scalable, and interpretable solution for mutation spectrum comparison that retains information from high mutation counts while accounting for biological replicate structure, and is freely available through the MutSeqR R package (Dodge et al., 2025) to support reproducible analysis across a broad range of spectral resolutions used in contemporary sequencing studies of mutagenesis.

## ACKNOWLEDGMENTS

We thank Sabit Cakmak and Erica O’Neill for reviewing the manuscript and providing constructive feedback during internal peer review.

## Data Availability

The Duplex Sequencing data used in this study are available from the Sequence Read Archive (SRA) under accession number PRJNA803048 or are available in the supplementary material of LeBlanc et al. (2022). The processed count matrices used for the examples are available within the MutSeqR package.

## Funding

This work was supported by the Genomics and Research Development Initiative to FM. CLY thanks the Canada Research Chairs Program.

## CONFLICT OF INTEREST STATEMENT

The authors declare that there is no conflict of interest.

## Supplemental Results

To characterize how the penalty affects the null distribution independently of any real spectral difference, we simulated both groups from a single shared Dirichlet-multinomial distribution matched to the observed Control-group spectrum (K = 6; Supplementary Figure 3). At λ = 0.001, the Hybrid and Fully Penalized null distributions were indistinguishable. At λ = 1, the Fully Penalized null had a mean about 48% lower than the Hybrid null, a shift toward zero with no replicates below zero in either implementation. A comparable but smaller shift (∼30%) appeared in the bootstrap null generated from the real BaP data. This null-level shift was consistently smaller than the penalty gap observed on real spectral differences (3- to 7-fold, at λ = 1), consistent with the analytical result that the gap scales with the squared distance between group spectra.

**Supplementary Table 1.**
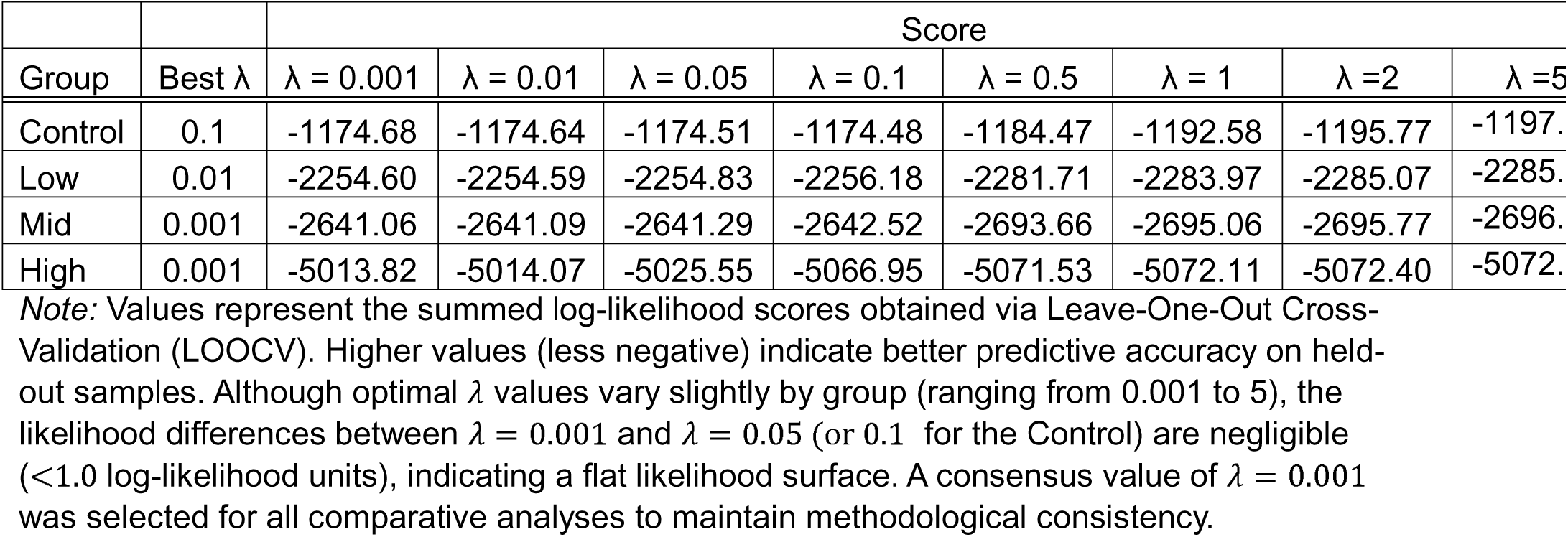
Optimization of the ridge regularization parameter (λ) via Leave-One-Out Cross-Validation.

Supplemental Table 2. Type I Error Rates (False Positive Rates) of the RP-DMM Across Varying Mutation Depths, Fold Differences, and Replicate Numbers in the 96-Channel Spectrum Context.

**Table S2 Note:** Type I error rates were evaluated across 1,000 simulated iterations under the true null hypothesis (identical underlying mutational spectra across groups). Simulations varied total mutation counts in the lower-mutational-burden group, biological replicate numbers (4-6 per group), and fold differences in mutation burden between groups. Expected mutations per category were calculated assuming a uniform distribution across the 96 mutation channels. Bold values highlight scenarios in which the observed false positive rate approached the nominal significance level (α = 0.05). Results indicate that reliable Type I error control was generally achieved when the lower-mutational-burden group contained approximately 300-500 total mutations, depending on the number of biological replicates, with approximately 400 mutations providing a practical and conservative recommendation for routine RP-DMM analyses.

**Supplemental Figure 1.**
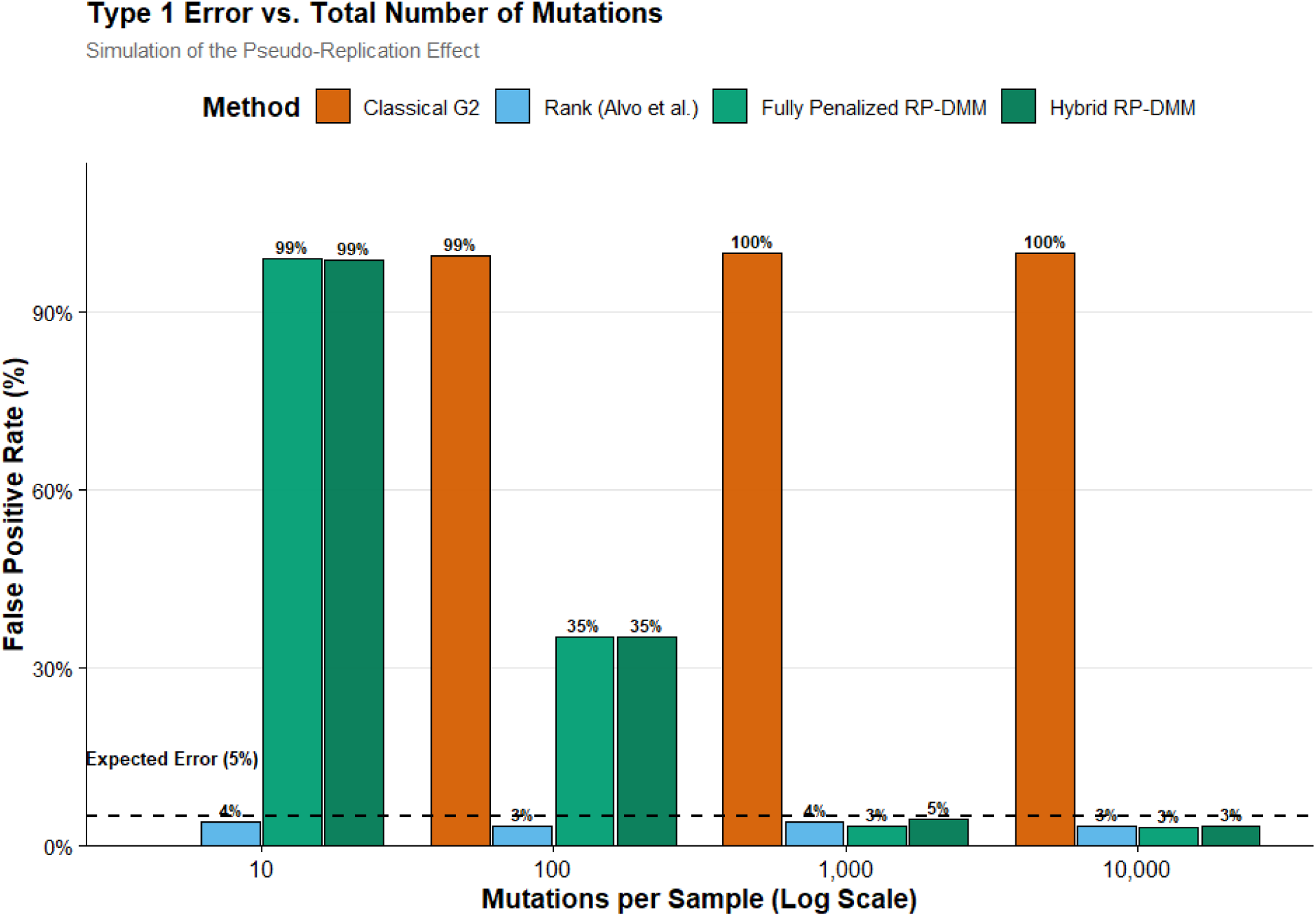
Type 1 error rates as a function of total mutation count for the 96-channel trinucleotide spectrum context (K = 96). At extremely low mutation depths (N = 10 per sample, group total = 40), both RP-DMM implementations exhibited severely inflated Type I error rates (99%), indicating insufficient information to reliably estimate the 96-category mutational spectrum and concentration parameter. At N = 100 mutations per sample (group total = 400), Type I error remained elevated under this specific simulation setting (four replicates per group), demonstrating that mutation counts only slightly exceeding the dimensionality of the spectrum are insufficient for stable inference. Reliable Type I error control (3-5%) was achieved at N = 1,000 mutations per sample (group total = 4,000) and above. These initial simulations suggested that substantially greater mutation depth is required for RP-DMM analyses of 96-channel spectra than for lower-dimensional spectral representations. Additional simulations using intermediate mutation depths and varying numbers of biological replicates (Supplemental Table 2) further refined this observation, indicating that reliable Type I error control was generally achieved when the lower-mutational-burden group contained approximately 300-500 total mutations, depending on replicate number, with approximately 400 mutations representing a practical target for routine applications. In contrast, the classical G² test exhibited near-nominal Type I error at N = 10 per sample (4%), primarily because the extremely sparse data limited its ability to detect any differences. As mutation depth increased, however, the G² test became progressively anti-conservative, reaching Type I error rates approaching 100%. The rank-permutation test maintained near-nominal Type I error rates (3-4%) across all tested depths, demonstrating robustness to both mutation depth and spectral dimensionality. Consequently, it may serve as a conservative alternative when mutation counts fall below those associated with reliable RP-DMM performance.

**Supplementary Figure 2.**
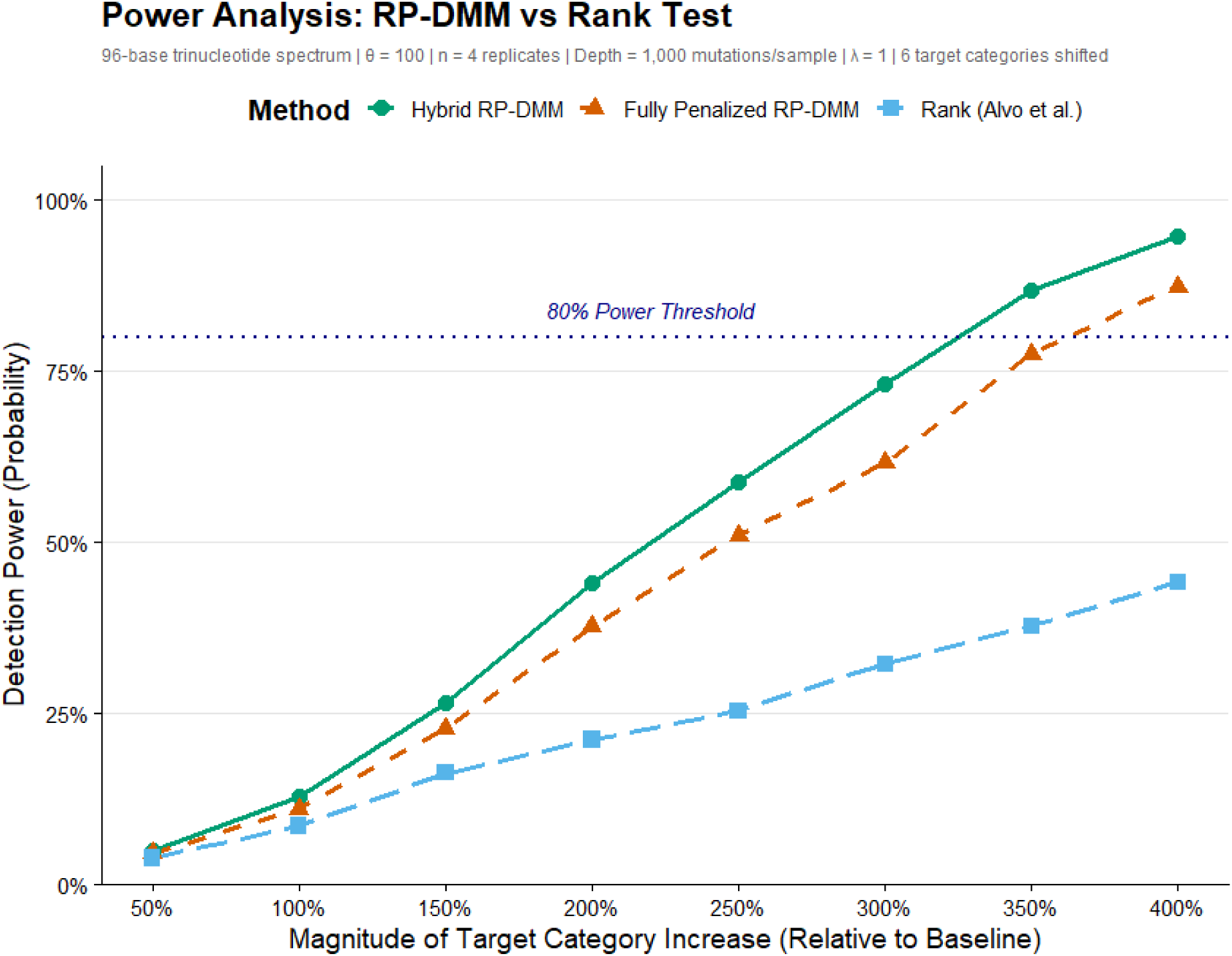
Power Comparison: Hybrid vs. Fully Penalized vs. Rank-based methods. Detection power comparison for the 96-channel trinucleotide spectrum context. A simulation study was performed to evaluate the sensitivity of the Hybrid RP-DMM (green), Fully Penalized RP-DMM (orange), and rank-permutation test (blue) in detecting a relative increase in a cluster of six target mutation categories. Simulations were conducted under realistic conditions: n = 4 replicates per group, 1,000 mutations per sample, moderate overdispersion (θ = 100), and regularization parameter λ = 1. Model parameters were jointly optimized under the ridge penalty for both RP-DMM implementations, with the Hybrid approach evaluating the constructed test statistic using the unpenalized likelihood (λ = 0) and the Fully Penalized approach evaluating it using the penalized likelihood (λ = 1). In the 96-channel context, where mutation counts are distributed across a large feature space and many categories are sparse, the Hybrid RP-DMM demonstrated meaningfully higher detection power than the Fully Penalized implementation across the entire tested range, crossing the 80% power threshold at approximately a 325% relative increase in the target categories compared to approximately 400% for the Fully Penalized approach. This divergence, which was not observed in the 6-channel context (Figure 2), reflects increased penalty-induced attenuation of the fully penalized test statistic in high-dimensional settings: the ridge penalty aggressively shrinks sparse category estimates toward the uniform baseline, and when this penalized likelihood is used for test statistic evaluation as well as parameter estimation, biological signals in sparse categories are partially suppressed. The Hybrid approach avoids this by decoupling stable parameter estimation from significance evaluation, allowing sparse category signals to contribute fully to the test statistic once the spectrum estimates have been regularized. The rank-permutation test exhibited the lowest detection power across all tested effect sizes, failing to reach 50% power even at a 400% relative increase, consistent with the known limitations of rank-based methods when count-magnitude information is discarded. These results indicate that the Hybrid evaluation strategy provides a meaningful power advantage over fully penalized evaluation in high-dimensional spectral analyses at larger regularization strengths (λ = 1); at the smaller, cross-validated λ used in the benzo[a]pyrene case study (λ = 0.001), the two implementations performed comparably (see Methods).

**Supplemental Figure 3.**
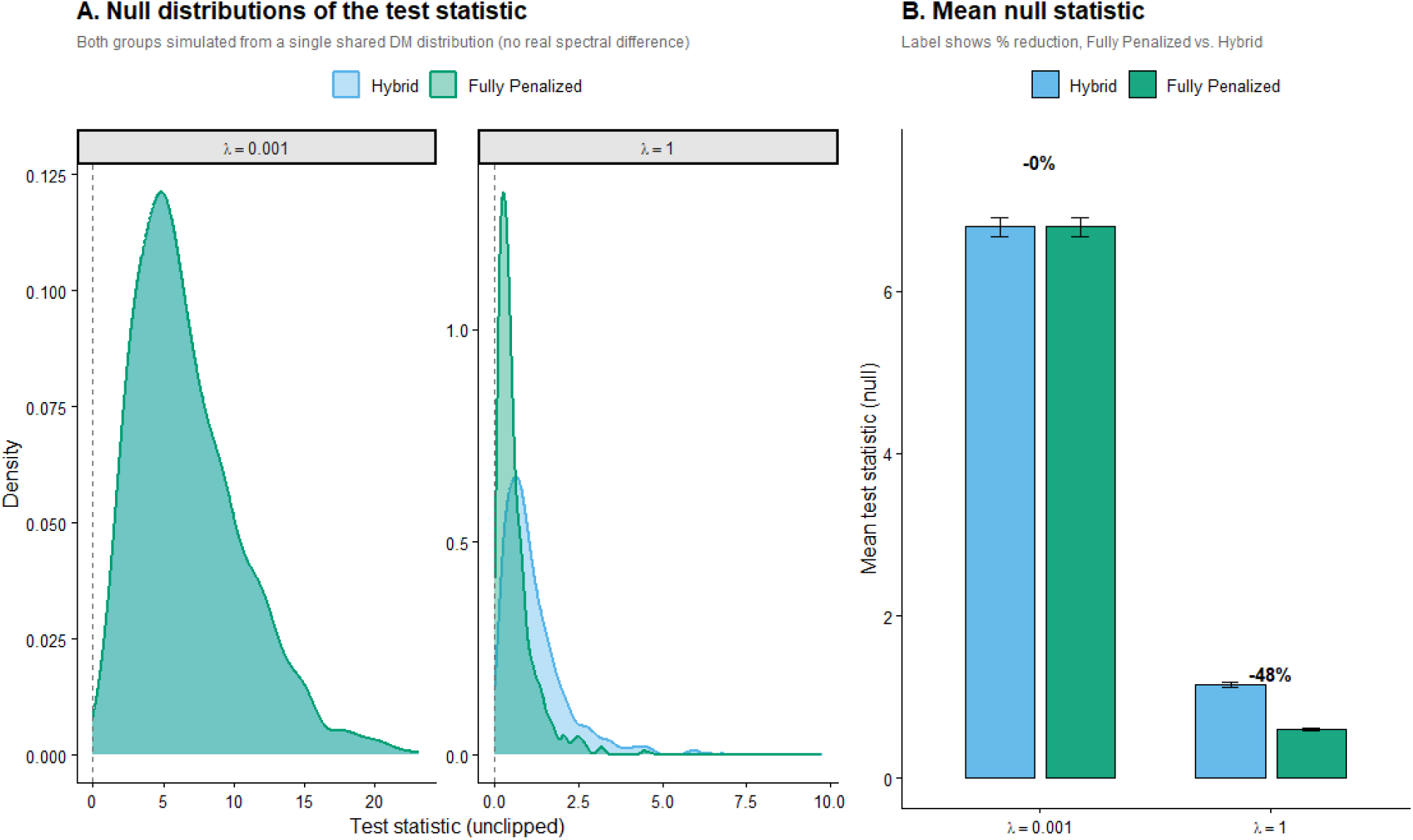
Effect of the ridge penalty on the null distribution of the Hybrid and Fully Penalized test statistics. To isolate the effect of the penalty from any real spectral difference between groups, both simulated groups were drawn from a single shared Dirichlet-multinomial distribution matched to the observed Control-group spectrum and sequencing depths (K = 6, 1,000 simulations per panel). (A) Density of the unclipped test statistic under each implementation, at λ = 0.001 and λ = 1. (B) Mean null statistic for each implementation at each λ (± SE), with the percentage reduction for the Fully Penalized statistic relative to Hybrid. At λ = 0.001, the two null distributions were indistinguishable (0% reduction). At λ = 1, the ridge penalty shifted the Fully Penalized null toward zero by 48%, with the density still visibly extending to zero in Panel A rather than piling up there, indicating a shift rather than a truncation. This null-level shift is smaller than the penalty gap observed between real, spectrally distinct groups (Table 3), consistent with the Fully Penalized statistic’s reduced power to detect true differences rather than any bias in its Type I error.

## Notes

https://www.ncbi.nlm.nih.gov/sra/?term=PRJNA803048

### Competing Interest Statement

The authors have declared no competing interest.

### Summary of Updates

Added Morrill Gavarro, L., Couturier, D.-L., & Markowetz, F. (2025). A Dirichlet-multinomial mixed model for determining differential abundance of mutational signatures. BMC Bioinformatics, 26, 59. https://doi.org/10.1186/s12859-025-06055-x reference Modified the introduction, methods and discussion as a result. Supplemental Table 1 was revised.

